# Using river temperature to optimize fish incubation metabolism and survival: a case for mechanistic models

**DOI:** 10.1101/257154

**Authors:** James J. Anderson

## Abstract

Allocating reservoir flows to societal and ecosystem needs under increasing demands for water and increasing variability in climate presents challenges to resource managers. In the past, managers have operated reservoirs to achieve flow and temperature compliance points based on qualitative predictions of competing needs. Because it is difficult, if not impossible, to assess whether meeting such compliance points is efficient or equitable, new strategies for regulation are being advanced. Critical to these strategies is the need for new models with sufficient biological details to identify the effects of reservoir operations on organism growth and survival in real time. This paper evaluates the adequacy of three models of differing complexity for managing the Sacramento River temperature during the incubation of winter-run Chinook salmon. The models similarly characterize temperature-and density-dependent mortality from egg through fry survival, but use different spatial and temporal resolutions. The models all fit survival data reasonably well, but predict different reservoir operations to protect fish. Importantly, the models with the finer spatial/temporal resolution predict reservoir operations that require less flow and better protect fish when water resources are limited. The paper illustrates that shifting the focus of management from meeting compliance points to meeting the metabolic needs of the organisms’ yields efficiencies and identifies when water is needed and when it can be saved.

## Introduction

With the construction of Shasta and Keswick dams on the Sacramento River in the 1940s the historical cold water upstream spawning habitat for the winter-run Chinook salmon (*Oncorhynchuys tshawytscha*) was blocked. The run now spawns in the warmer reaches between Keswick Dam and Bend Bridge (Fig.1) and with this displacement out of its natural habitat the run was judged to be at a low to moderate risk of extinction (Lindley et al. 2007). In particular, the run was considered vulnerable to catastrophes such as a prolonged drought that depletes the cold water storage of Shasta Reservoir or some failure of cold water management (Williams 2006). These possibilities, predicted in 2006, were realized with the onset of California’s longest drought beginning 2011 (Seager et al. 2015) and the inability of the Shasta Reservoir operations to access the diminished cold water from the bottom of the reservoir in 2014. As a result, for three consecutive spawning seasons (2013-2015) the incubating winter-run eggs and alevin experienced unprecedented warm waters that resulted in historically low survival.

**Figure 1.**
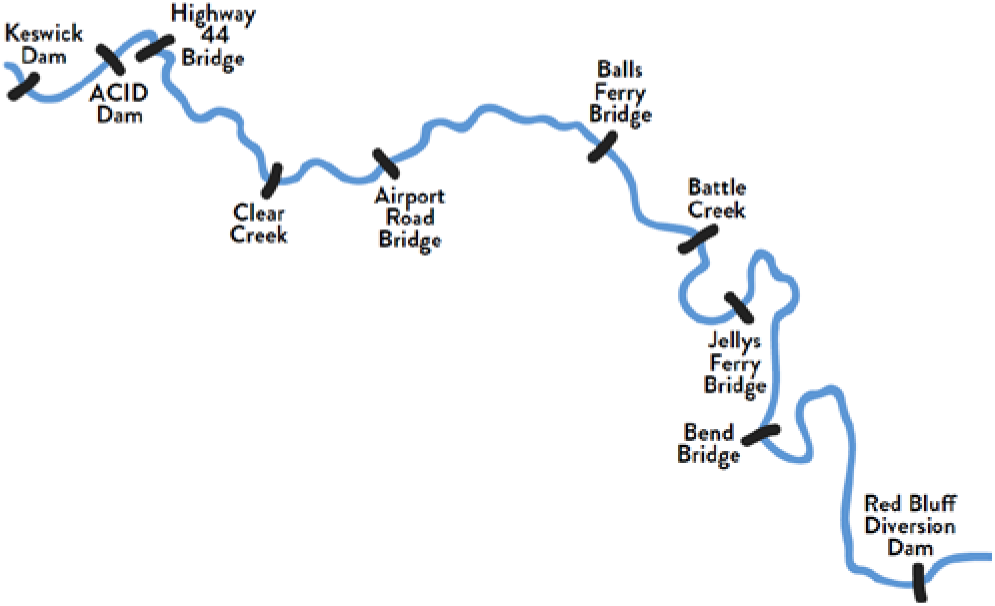
Upper Sacramento River showing reaches of winter-run Chinook salmon spawning habitat.

Recent modeling, on the effects of temperature on survival from egg deposition to fry passage at Red Bluff Diversion Dam (RBDD), suggested that the historically low survivals were indeed the result of high incubation temperatures (Martin et al. 2017). Based on firsts principles of egg metabolism and oxygen flux in redds the authors concluded that the lethal incubation temperature for eggs in the river was 3 °C lower than that determined in laboratory studies. Following on the new model, the National Marine Fisheries Service (NMFS) proposed amendment to the Biological Opinion on the long-term operations of the Central Valley water projects that better links the Sacramento River temperature control program to the biological metrics of winter-run Chinook spawning and egg incubation (NMFS 2017). The proposed regulations specify that between May 15 and October 31 the Bureau of Reclamation operate Shasta Reservoir to maintain the temperature between Keswick Dam and the confluence with Clear Creek (Fig. 1) at or below a compliance temperature point determined by the model. Also, because the ability to regulate temperature depends on the cold water resources in the reservoir, the compliance temperature would be adjusted according to the cold water resource with compliance set at 53 °F (11.7 °C) in normal and above water years, at 54 °F (12.2 °C) in dry years and 56 °F (13.3 °C) in critically dry years.

The proposed regulation for Shasta Reservoir is an important example of the shift to biologically-based management that seeks to balance the increasing demands of limited water resources for ecosystem services and human consumption (Poff et al. 2015, Poff and Schmidt 2016). A study of Mekong River management (Sabo et al. 2017) to promote both fish and hydroelectric production succinctly framed this shifting paradigm from addressing “How much water do we need?” to “When do we need it most and when can we spare it?” Many of the studies promoting this new paradigm link management actions to biological responses through statistical correlations. The NMFS example is unique in that the proposed regulation is based on a mechanistic model. The question thus arises, does the model describe the biological mechanisms with sufficient resolution to identify reservoir operations that efficiently and adequately protect fish? To address this question this paper compares the egg mortality model of (Martin et al. 2017) with extended versions that include additional information and finer resolution of the biological processes.

## Methods

The strategy formulates the survival of winter-run Chinook salmon from egg fertilization in the reaches below Keswick Dam to passage of fry at RBDD using three different levels of physiological and spatial resolution. The three models, based on the framework of (Martin et al. 2017), include temperature-dependent and density-dependent mortalities, denoted here thermal and background mortalities. Model I, developed by (Martin et al. 2017), expresses thermal mortality independent of the stage or age of incubation and background mortality independent of redd location. Model II extends model I with age-dependent thermal mortality and model III adds river reach-specific background mortality.

### Model I: age-independent thermal mortality, spatially-independent background mortality

In Martin et al. (2017), survival of fertilized eggs, deposited in redds below Keswick Dam to their passage at RBDD as fry, is described with temperature-and density-dependent mortality processes. The daily pattern of temperature, generated from a high-resolution river temperature model, is used to determine the daily mortality rate and fry emergence date for each redd. The background mortality is defined at the coarser resolution of the total number of spawners in the habitat. Importantly, the annual egg-to-fry survival depends on thermal mortality independent of the stage of incubation and background mortality independent of the redd location. The relationship of total survival with the driving covariates can be expressed for each cohort as

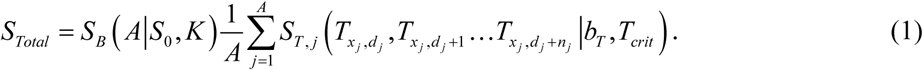

Background survival *S*_*B*_ depends on total number of spawners *A* in the habitat in the year and is defined with a Beverton-Holt model (Beverton and Holt 2012) as

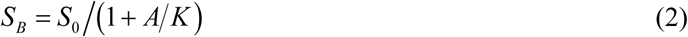

where the parameter *S*_0_ is the maximum background survival when adults do not compete for redd space and *K* is the carrying capacity. The Beverton-Holt model describes the incremental effects of increasing spawner abundance on juvenile survival. Both parameters are assumed constant across years.

Thermal survival *S*_*T*, *j*_ defined by eq. (3) calculates survival for each redd, *j*, using its location *x*_*j*_ and day of year (DOY) of formation *d*_*j*_, which then defines the sequence of temperatures *T*_*x*_*j*_, *d*_*j*__, *T*_*x*_*j*_, *d*_*j*_+1_ … *T*_*x*_*j*_, *d*_*j*_+*n*_*j*__ experienced by fish in the redd between *d*_*j*_ and DOY of fry emergence, *d*_*j*_ + *n*_*j*_. The daily survival rate depends on the differential between the daily *T*_*x*_*j*_, *d*_*j*_+1_ and the critical *T*_*crit*_. Mortality only occurs on days that the differential is positive. The thermal survival in redd *j* over its incubation duration is then

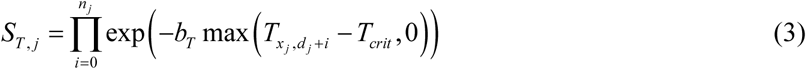

where *b*_*T*_ fixes the mortality rate per degree of temperature differential. The duration *n*_*j*_ of incubation from egg fertilization to fry emergence depends on temperature according to (Zeug et al. 2012).

### Model II: age-dependent thermal mortality

Model II includes age-dependent thermal mortality. To accommodate the added complexity, the model represents the population in terms of the mean thermal conditions of incubation in five spawning reaches below Keswick Dam (Table 1). Additionally, the model assumes mortality occurs only at a critical age of development *a*_*crit*_. The critical DOY *d*_*crit*_ and critical age are represented by lagging back in time the mean emergence DOY *d*_*emerge*_ and mean emergence age *a*_*emerge*_ by *δ* days giving

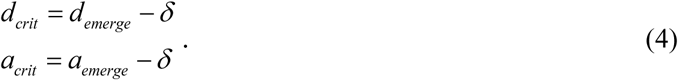

**Table 1.** River reach length, *L* and spawning reach index, *i*.

| $i$ | River Reach | $L_i$ (km) |
| --- | --- | --- |
| 1 | Keswick Dam to A.C.I.D. Dam | 5.8 |
| 2 | A.C.I.D. Dam to Highway 44 Bridge | 3.4 |
| 3 | Highway 44 Bridge to Airport Road Bridge | 19.3 |
| 4 | Below Clear Creek to Balls Ferry Bridge | 12.2 |
| 5 | Balls Ferry Bridge to Bend Bridge | 30 |
|  | Keswick Dam to Red Bluff Diversion Dam | 94 |

The mean age of emergence is determined from the cumulative thermal exposure according to (Zeug et al. 2012). The variable *δ* is a free parameter that can assume any value between 0 and *a*_*emerge*_. For simplicity, terms in eq. (4) are applied for all reaches in a year but vary annually. The model framework similar to model I is expressed

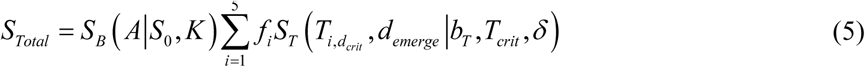

where *ƒ*_*i*_ is the fraction of the total female spawning population in reach *i*. The differential temperature on the critical date corresponding to *δ* is *T*_*i*, *d*_*crit*__ - *T*_*crit*_ and thermal survival is

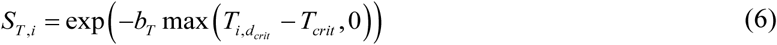

where *T*_*crit*_ and *b*_*T*_ are regression parameters.

Model II only allows mortality at a single critical DOY and age, which are characterized by the lag (4). In reality, temperature-dependent mortality might occur on any day, as is assumed in model I, but studies considered in the discussion suggest mortality occurs in a limited range of days about a mean critical ag *ā*_*crit*_ or over several distinct developmental stages, e.g. egg hatch and maximum total dry mass (MTDM) (Rombough 1994, Rombough 2007). In effect, Model II provides a simplified way to explore the possible ages of thermal sensitivity in terms of *δ*. Candidate values of *δ* are required to yield good fits to data and critical ages corresponding with metabolically critical stages.

### Model III: age-dependent thermal mortality, spatially-dependent background mortality

Model III expands on model II by expressing reach-specific background mortalities. The framework is

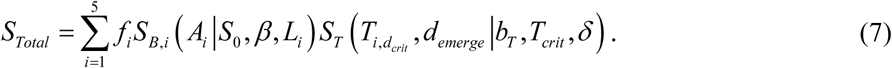

Each reach has a carrying capacity defined by its length *L*_*i*_ as *K*_*i*_ = *βL*_*i*_ where *i* specifies spawning reaches *i* = 1, 2, 3, 4, 5 (Table 1) and *β* is a constant with dimensions of redds/km characterizing carrying capacity per unit length. Background survival is then

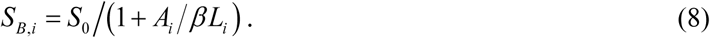

### Model differences

The three models apply equivalent thermal and background mortality rate equations but differ in their spatial/temporal representations of the covariates. Model I defines the location-specific temperature experience of individual redds on a daily basis. It thus characterizes the incubation duration for individual redds. Models II and III use a reach-specific spatial scale and track the daily running average of temperature for the average redd in each reach. The incubation duration in eq. (4) is based on the average incubation duration in reach 1. While the three models characterize mortality in terms of the difference between the environmental temperature and a critical temperature, the age-dependency of the rates differ. Model I allows mortality for any incubation age at which the differential is positive, whereas models II and III only allow mortality at a specified critical age of development. Finally, the spatial scale of background mortality differs; models I and II characterize background mortality at the river scale and model III characterizes at the reach scale.

### Model fitting

All models were fit to logit-transformed egg-to-fry survival data. For model I parameters *S*_0_, *K*, *b*_*T*_ and *T*_*crit*_ were obtained from (Martin et al. 2017). For models II and III *d*_*emerge*_ and *a*_*emerge*_ for were estimated from the SacPAS model (www.cbr.washington.edu/sacramento/migration/), which uses the observed spawning dates and temperatures referenced to Keswick Dam to calculate the emergence date from each redd according to (Zeug et al. 2012). The temperature *T*_*i*, *d*_*crit*__ in reach *i*, corresponding to a critical DOY, was expressed as a 6-day running average. Using the *optim* function in R^©^ (R Core Team 2016) the parameters defining *S*_*Total*_ in models II and III were estimated by minimizing ∑(log(*S*_*obs*_/(1 - *S*_*obs*_)) - log(*S*_*Total*_/(1 - *S*_*Total*_)))^2^. Coefficients for model II (*S*_0_, *K*, *b*_*T*_ and *T*_*crit*_) and model III (*S*_0_, *β*, *b*_*T*_ and *T*_*crit*_) were estimated for trial lags from *δ* = 1 to 80 d. The age of emergence varied annually over the range 75 < *a*_*emerge*_ < 90 with a representative value taken as *a*_*emerge*_ = 80 d. In essence, using the mean population emergence day, temperature data and egg-to-RDDD survival, the algorithm estimates the parameters and r-squares for each trial value of *δ*. Values of *δ* were then judged according their r-squares and the biological significance of the corresponding *a*_*crit*_.

### Data

The data for years 2002-2015 were fit to the model. Temperature data was derived from the SacPAS data site http://www.cbr.washington.edu/sacramento/data/, which has data downloaded from the California Data Exchange Center http://cdec.water.ca.gov/. Temperatures at midpoints of reaches were estimated by linear extrapolation between Keswick Dam and Balls Ferry Bridge from temperature monitors. Redd count data was obtained from California Department of Fish and Game aerial redd surveys (2017 Aerial redd Sacramento River summary as of 9-27-2017, from Doug Kilam) Mean emergence dates were generated with SacPAS. Profiles of weekly spawning numbers and predicted emergence dates and daily temperatures over years 2001-2015 are given in Supporting Information Fig. S1. Survival from egg fertilization to fry passage at RBDD was reported by CDFG and extracted from (Martin et al. 2017). For comparison with carcass counts, the redds were regressed against carcass counts obtained from (Poytress 2016) yielding one redd equates to 5.7 carcasses with r-sqr = 0.85.

## Results

The analysis focuses on model III but results of model I and II are provided for comparison. The mortalities in model III are driven by reach temperatures and observed redd depositions. To set the models in context, first consider the temperature and fish patterns used in the models.

### Temperature and spawner distributions

Figure 2a and 2b illustrate the temperatures in reaches at spawning and near egg hatching. In most years temperatures in reaches 1 and 2 were below 12 °C while the temperatures in reach 4 were near or above 12 °C. Figure 2c illustrates the fraction of the total population in each reach. In most years the greatest fractions were in reaches 1 and 2. The spawning density, expressed as females/km, varied significantly across reaches and were generally highest in reach 1 and 2 (Fig. 2d). Reaches 4 and 5 each had about 0.3% of the total spawners. Note because of its greater length, reach 3 had low density (Fig. 2d) but contained appreciable fractions of the population (Fig. 2c). The relative fractions of fish between reaches is illustrated in Fig. 3. Note the fractions in reach 1 and 2 were inversely correlated while the fractions in reaches 2 and 3 were uncorrelated. The partition of redds amongst the reaches was in part affected by the pre-spawning water temperature (Fig. 4). In particular, when the pre-spawning temperature in reach 4 was > 12 °C the fraction of redds in reach 3 was below-average (Fig. 4c). However, the distributions in reaches 1 (Fig. 4a) and 2 (Fig. 4b) were uncorrelated with the pre-spawning temperature.

**Figure 2.**
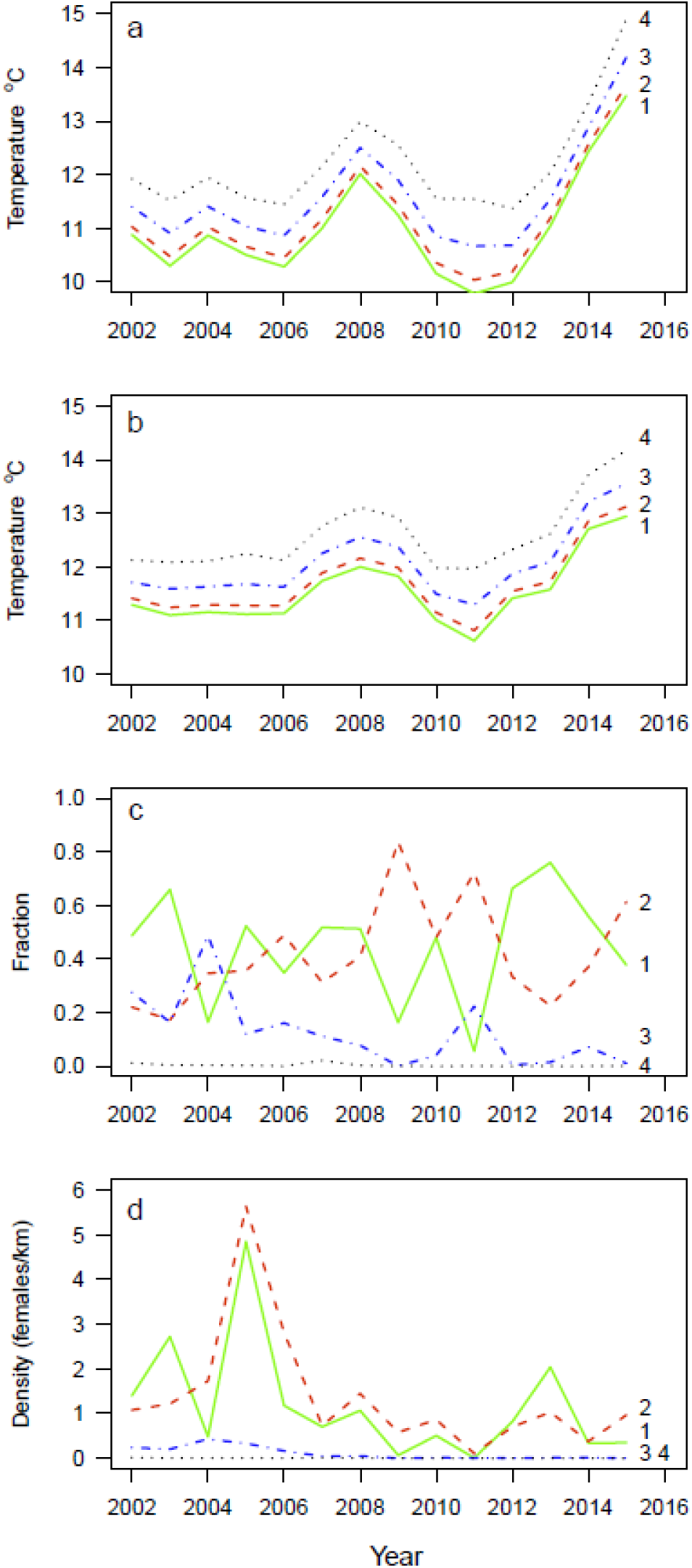
Reach temperature prior to spawning (a), at mean hatch DOY (~ 37 d) (b), fraction of redds in reach (c), and density of redds in reach (d). See Table 1 for river reach specifics.

**Figure 3.**
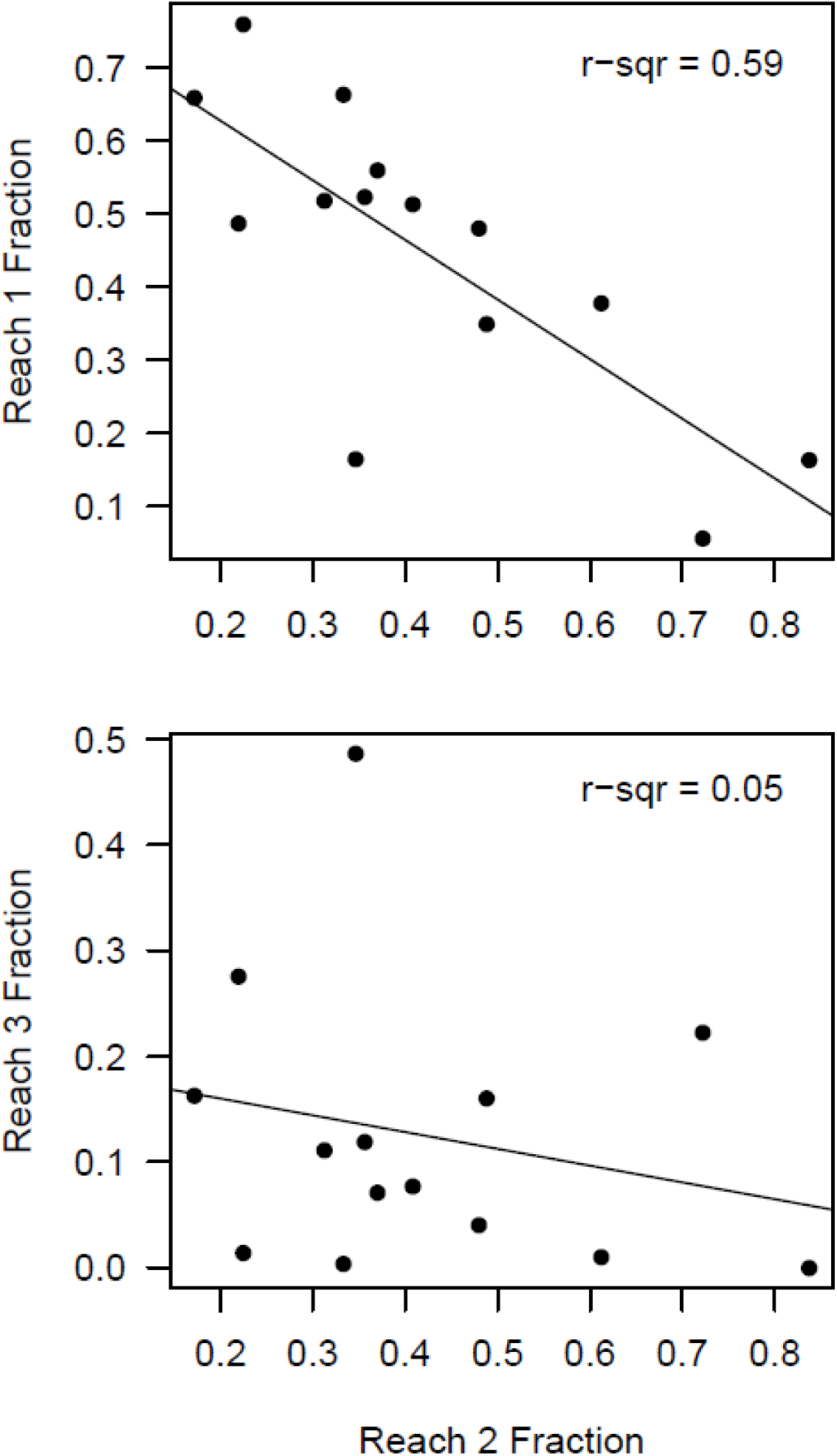
Relationship of fractions of population in reach 2 relative to reach 1 and reach 3.

**Figure 4.**
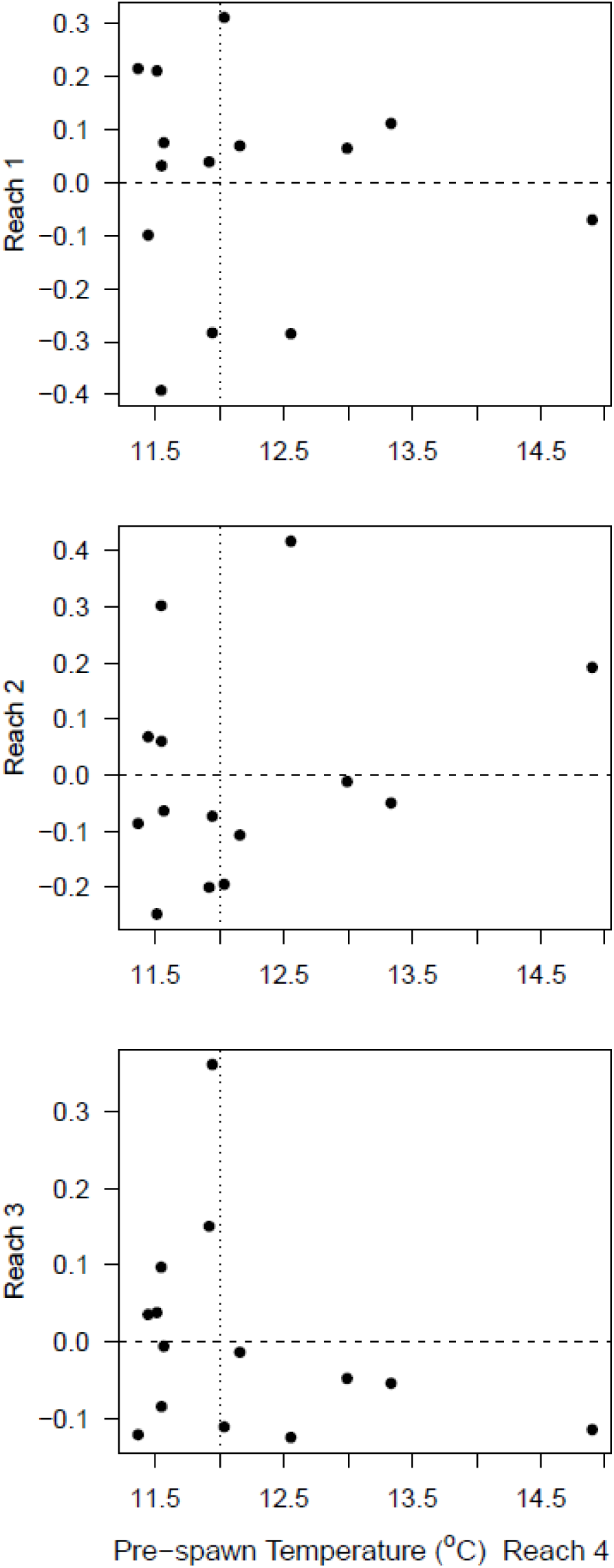
Deviations from mean fraction of redds in each reach plotted against the pre-spawning temperature in reach 4.

### Model fits

Models II and III were fit with all possible ages of critical thermal sensitivity, *a*_*crit*_ (Fig. 5). Notably both models produce high r-square fits for *a*_*crit*_ within the ages of egg hatching, approximately 40 d (Rombough 1994). Model III also produced high r-squares at *a*_*crit*_ = 24 and 67 d. The first peak has no clear biological basis, but the peak at 67 d corresponds with the age of alevin MTDM, which is also the maximum metabolic rate and occurs just prior to fry emergence (Rombough 1994).

**Figure 5.**
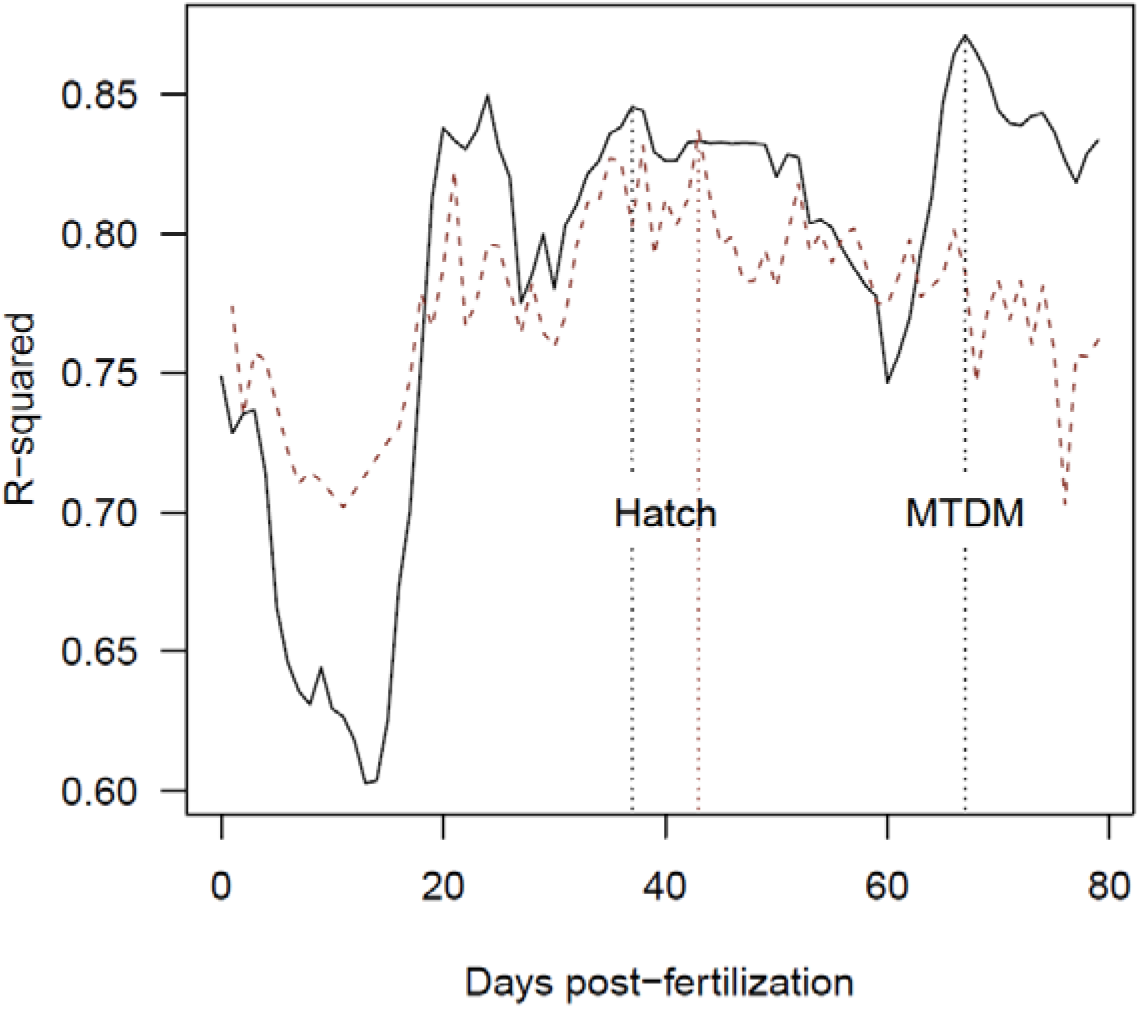
R-square generated from model fits vs. age of critical thermal sensitivity. Vertical lines at age 37 and 67 d correspond with approximate ages of egg hatch and MTDM. Model II results (---) and model III results (—).

The patterns of survival across years for the three models are illustrated in Fig. 6. For model III, a critical age corresponding to MTDM fits data slightly better than the age corresponding to egg hatching. Model II, with critical age at hatching, has a similar fit while model I underestimates the higher survivals. Judged in terms of r-square, model III and II best fit the data. (Table 2).

**Figure 6.**
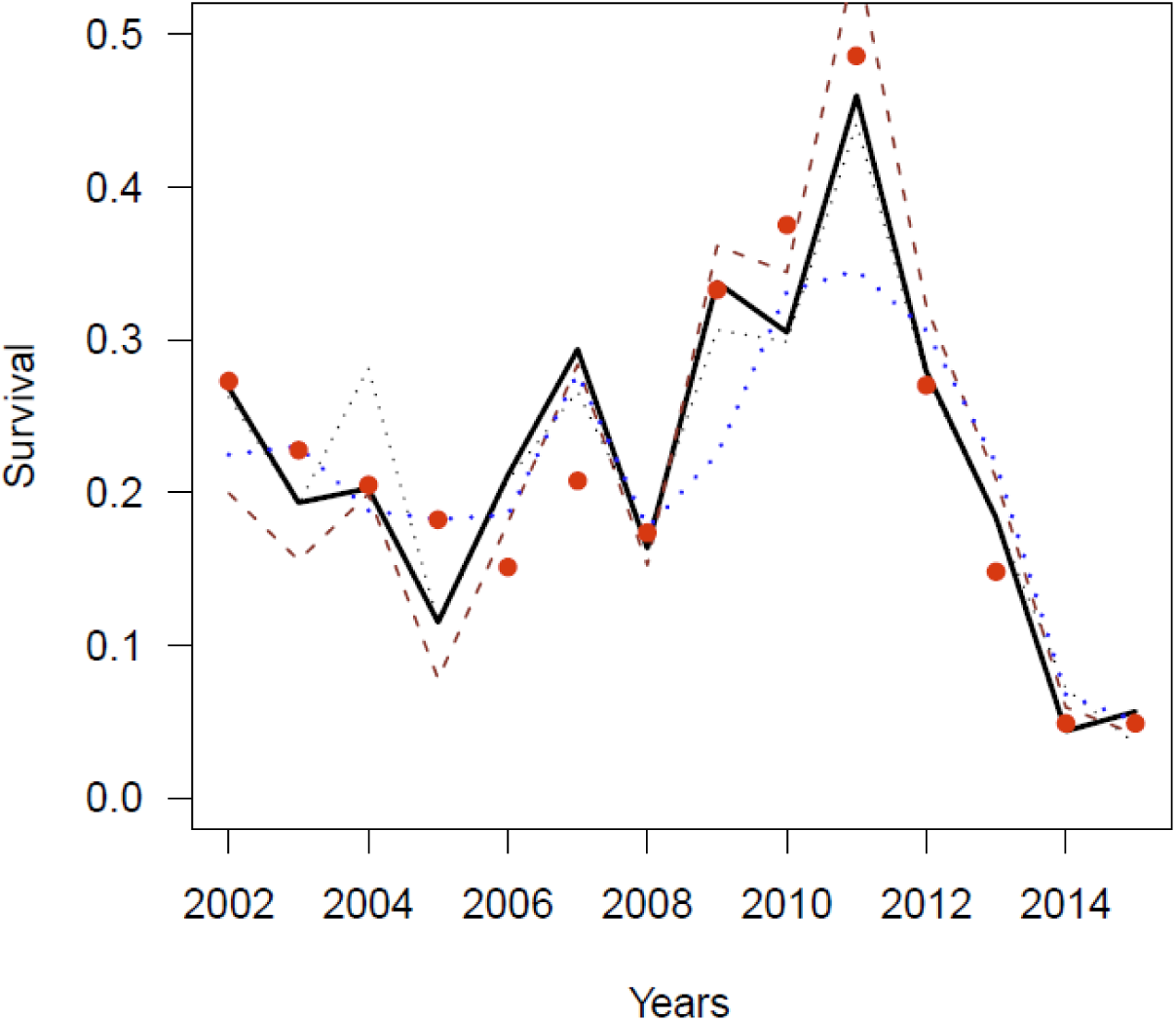
Comparison of survivals for model I (---), model II with critical age at hatch (--) and III with critical age at hatch (·····) and MTDM (—).

**Table 2.** Model parameters.

| Model | $a_{crit}$<br>(d) | $S_0$ | $T_{crit}$<br>(°C) | $b_T$<br>(d<sup>-1</sup> °C<sup>-1</sup>) | $\beta$<br>(redd km<sup>-1</sup>) | $K$<br>(carcass) | $r-sqr$ |
| --- | --- | --- | --- | --- | --- | --- | --- |
| I | - | 0.37<sup>a</sup> | 12.0<sup>a</sup> | 0.024<sup>a</sup> | - | 9107<sup>a</sup> | 0.77<sup>a</sup> |
| II | 43 | 0.58 | 11.8 | 1.92 | - | 1842<sup>b</sup> | 0.84 |
| III | 67 | 0.49 | 12.2 | 2.44 | 36.6 | - | 0.87 |
| III | 37 | 0.47 | 11.9 | 1.91 | 39.1 | - | 0.85 |

a. obtained from (Martin et al. 2017)
b. carrying capacity in redds and expanded to carcasses using 5.7 carcasses per redd

The patterns of model III parameters vs. critical age are illustrated in Fig. 7. The figure also demarks critical ages associated with egg hatching and fry MTDM and the highest r-squares (Fig. 5). Therefore, parameters for other critical ages are not supported by statistical or biological criteria. Noteworthy, the critical temperature increases with increasing critical ages of thermal sensitivity (Fig. 7c). Also, the maximum background survival *S*_0_ (Fig. 7a) and the carrying capacity parameter *β* (Fig. 7b) vary inversely such that high estimates of *S*_0_ correspond with lower estimates of carrying capacity. Thus, the observed pattern of background survival can be achieved with a range of correlated background parameters.

**Figure 7.**
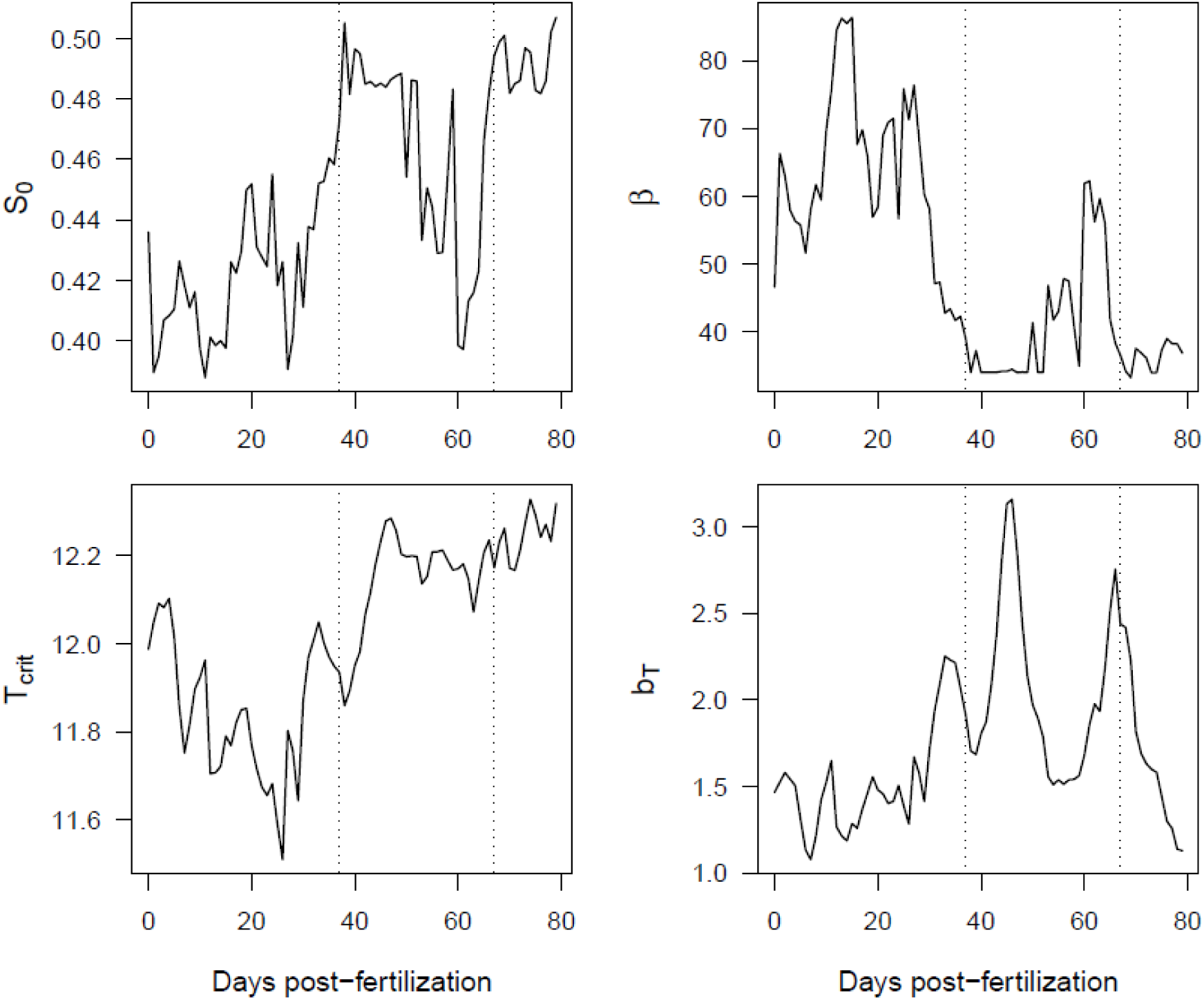
Model parameters as a function of possible ages of thermal sensitivity *a*_*crit*_. Vertical lines at post-fertilization day 37 (·····) and 67 (·····) correspond with approximate ages of egg hatch and MTDM.

Parameters for the models are given in Table 2. Model I parameters are extracted from (Martin et al. 2017). Model II parameters are shown for critical ages 43 d, which corresponds to the high r-square and is the approximate age of egg hatching. Model III parameters for critical ages 37 and 67 d, which have high r-squares, are candidates for ages of hatching and MTDM (Fig. 5). Notably, the maximum background survival, *S*_0_, in models I and III are different but the critical temperatures are similar. The mortality rate term, *b*_*T*_, is a factor of 100 lower in model I than in models II and III because mortality is spread over several months in model I while it is restricted to the day of the critical age in models II and III. Models I and II estimate carrying capacity for the entire habitat while model III estimates capacity for each reach. The total carrying capacity for model III was derived as the sum of capacities for reaches 1 through 4. Reach 5 was excluded because it only contained 0.03% of the spawners. Note, the habitat carrying capacities for models I and III are equivalent.

Model III reach-specific thermal and background survivals are illustrated in Fig. 8. The intensity of the mortality processes in the reaches vary inversely with the survival. Figures 8a and 8b illustrate the reach survivals under assumptions that the critical age for thermal mortality are at hatching and MTDM. Note that the distribution of mortalities depends on the critical age. The survival patterns for reaches 1 and 2 are similar for both critical ages but the patterns of survival for the lower two reaches varies with the assumption of critical age. The model indicates higher mortality would occur in the lower reaches for fish susceptible to thermal mortality at the alevin pre-emergent age. This pattern results because MTDM occurs approximately 30 days after the hatch and therefore fish are exposed to higher temperatures at the day of MTDM relative to the day of hatching. Background mortality is essentially independent of the critical age assumption (Figs. 8c). Background survival increases from reach 1 to 4, which is essentially opposite the pattern for thermal survival.

**Figure 8.**
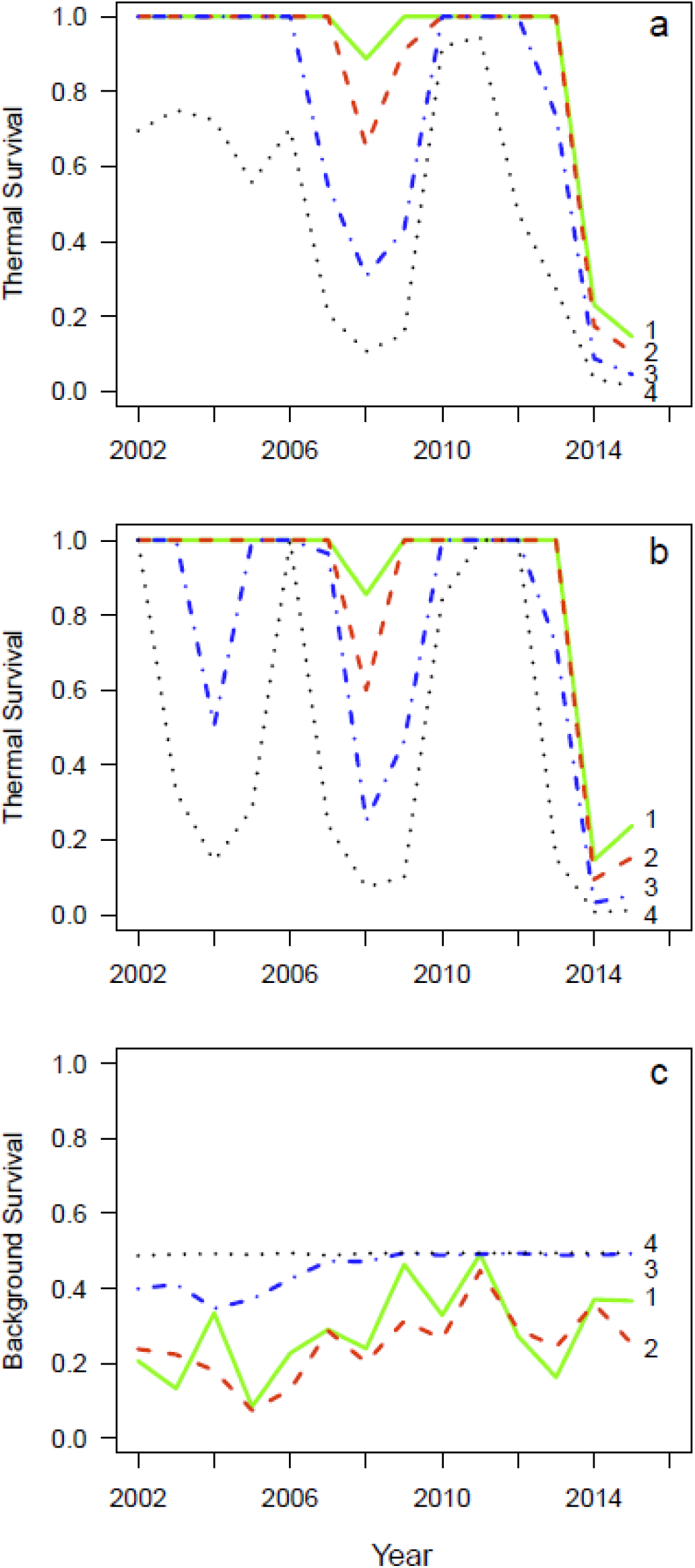
Thermal mortality in reaches 1-4 for critical day at hatch (a) and at MTDM (b). Background mortality for reaches 1-4 for critical day at either hatch or MTDM (c).

## Discussion

The three models use similar thermal and background mortality processes and produce similar fits to the annual patterns of egg-to-fry survival. However, the different spatial/temporal scales of the processes have significant implications for developing temperature management programs for the upper Sacramento River. Thus, support for the alternative models is discussed below.

### Age-dependent mortality

The three models characterize the thermal mortality rate in terms of the differential between the environmental temperature and a critical temperature as proposed by (Martin et al. 2017). In all models the best estimate of critical temperature is ~12 °C (Table 2). However, the models differ in the duration of mortality. In model I the mortality occurs any day the temperature differential is positive. In models II and III, incubating fish are only susceptible to mortality at the critical age. These two alternatives represent the extremes of possibilities and result in different predictions for reservoir operations. Fortunately, biological information is available on which to compare the alternatives.

For this comparison, first consider Fig 9 redrawn from (Rombough 1994) showing that over the incubation, the Chinook salmon egg/alevin metabolic demand per individual increases in an exponential-like manner between fertilization and the MTDM. Hatching is an important transition in the incubation because eggs obtain oxygen by diffusive flux across the egg membrane while alevin obtain oxygen by pumping water through their gills (Wells and Pinder 1996). This switch from diffusive to pump transport, greatly increases the fish’s ability to tolerate low oxygen after hatching. Figure 10 illustrates the transition and increased tolerance in terms of *P*_*c*_, the critical oxygen level below which routine metabolism becomes dependent on the ambient oxygen (Rombough 2007). *P*_*c*_ rises steadily during egg development and then drops about 50% after hatching. Additionally, the post-hatching oxygen sensitivity is essentially independent of incubation temperature (Rombough 1986). Thus, sensitivity to low oxygen increases steadily during egg development and then drops rapidly and stabilizes in the alevin stage. In essence, this transition in oxygen sensitivity is important because organisms with insufficient oxygen are susceptible to mortality and the eggs are significantly more susceptible than alevins.

**Figure 9.**
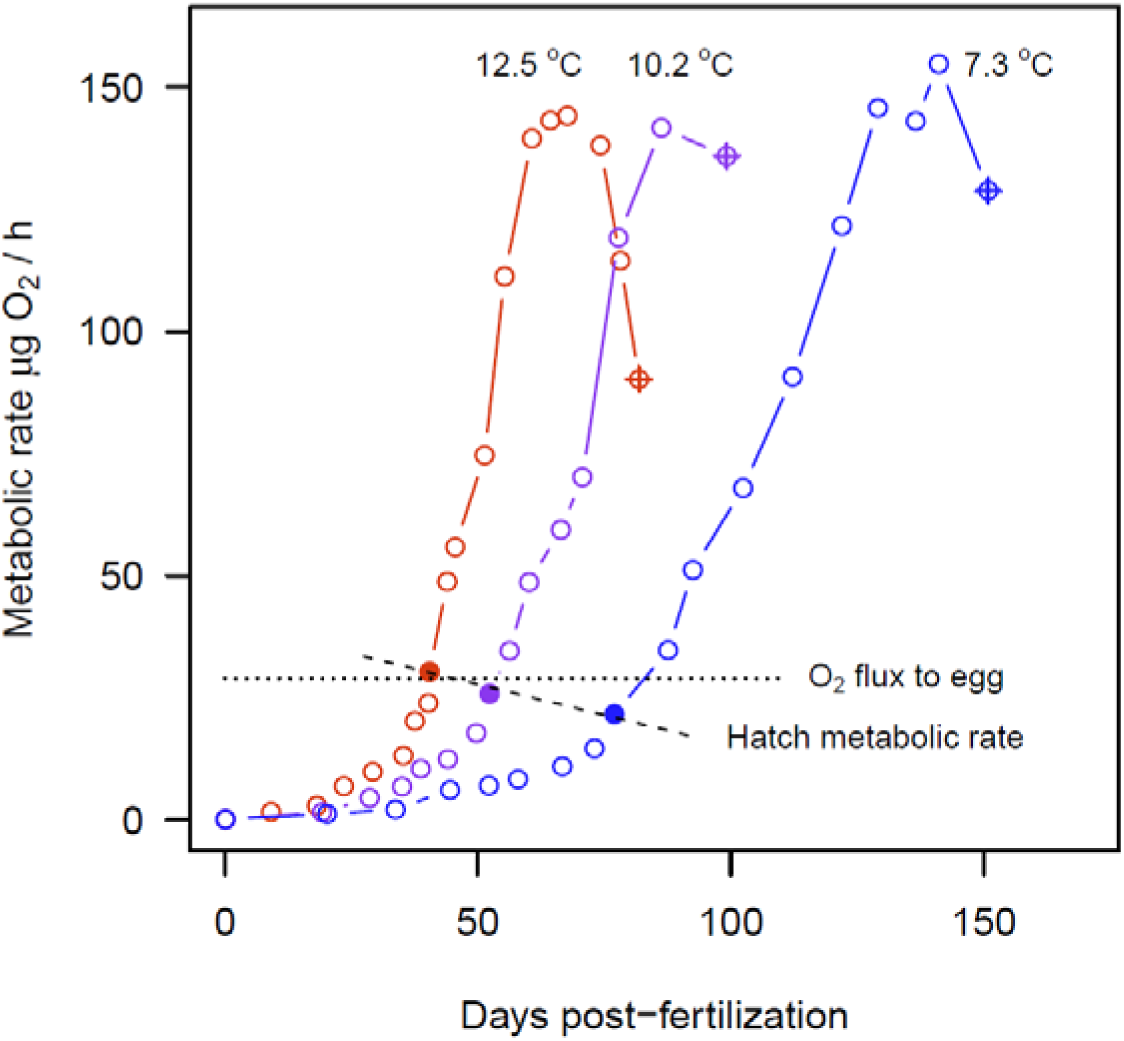
Incubating Chinook salmon metabolic rate vs. days post-fertilization at three temperatures. Redrawn from (Rombough 1994). Dotted lines depict hypothetical oxygen flux to embryo in redd required to induce mortality at temperature > 12 °C. Dashed line characterizes increases in hatch metabolic rate with temperature. The metabolic rate at hatching denoted (•), the peak metabolic rate corresponds to the Maximum Total Dry Mass (MTDM) and the rate at alevin emergence denoted (⊕).

**Figure 10.**
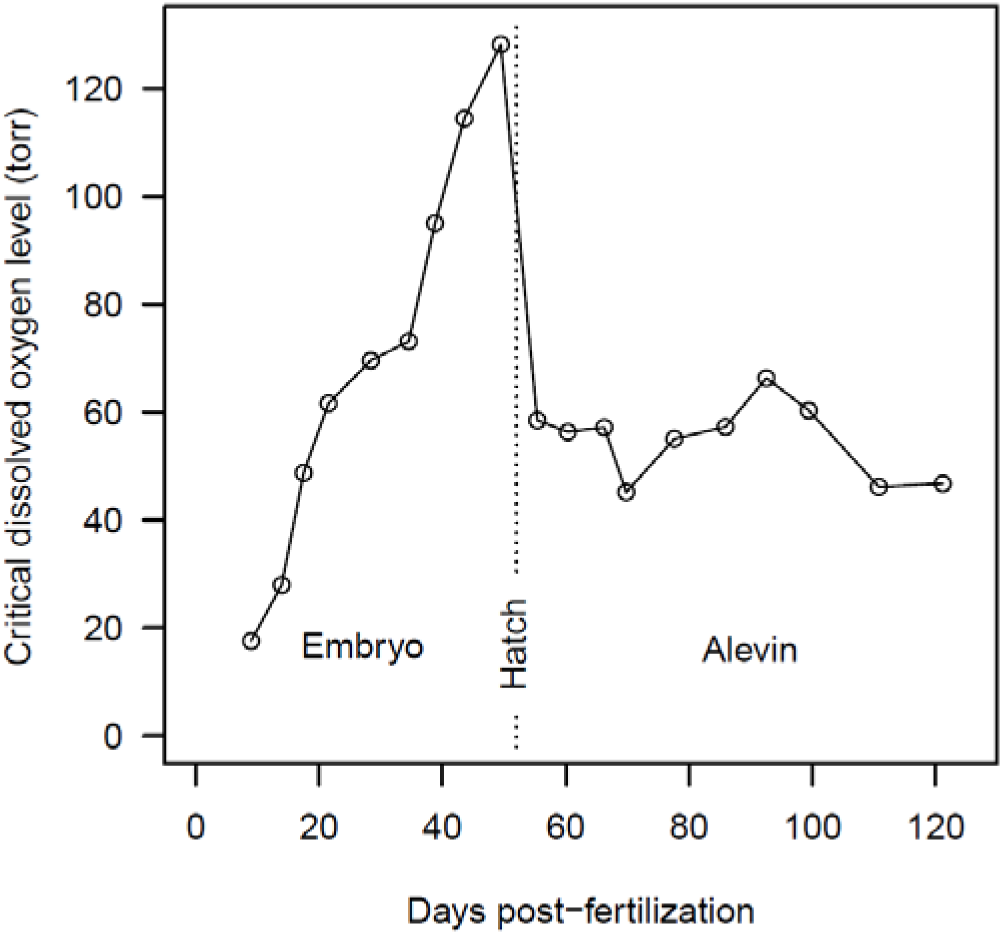
Critical oxygen as function of post-fertilization days for Chinook embryos and alevins at 10.2 °C and initial egg size 341 mg. Redrawn from (Rombough 1986).

Second, consider the model-derived critical temperature in the context of the changing metabolic rate. To illustrate, envision the fates of two groups incubating in redds in the Sacramento River at different temperatures. A blue fish group, incubating at a temperature of 10.2 °C follows the corresponding metabolic trajectory in Fig. 9 and at hatching has a metabolic demand of 26 μgO_2_/h. A red fish group, incubating at a temperature of 12.5 °C follows its corresponding trajectory and at hatching has a demand is 30 μgO_2_/h. In the alevin stage the metabolic rates of both groups continue to follow their respective trajectories but Fig. 10 indicates that their critical oxygen sensitivities after hatching both drop by half. Thus, because the red fish group temperature is above *T*_*crit*_ it experiences mortality and therefore prior to reaching the alevin stage its oxygen demand exceeds the diffusive oxygen flux available. Correspondingly the blue fish group experiences no mortality because its oxygen demand prior to hatching is below the diffusive flux. It is reasonable to conclude that the diffusive oxygen flux in redds lays between the metabolic rates of the two groups at hatching, i.e., ~28 μgO_2_/h. Additionally, the rate of increase in metabolic demand just prior to hatch increases at approximately 3 μgO_2_/h per day. Thus, the duration of low oxygen vulnerability is expected to occur just over a few days around hatching since a few days before hatching the egg oxygen demand would be significantly below the maximum diffusive flux and after hatching alevin obtain oxygen through gills.

Model III with a MTDM critical age (Table 2, Fig. 5) also suggests respiratory stress and associated thermal mortality could occur near the MTDM. Here laboratory support for later stage mortality is inconclusive. In Fig. 9 the maximum metabolic rate was independent of incubation temperature for Chinook salmon. If this were the case in the field environment, then thermal mortality at MTDM could occur at all incubation temperatures. The three models cannot resolve this possibility because such mortality would be attributed to background mortality, independent of temperature. However, other studies find a clear and well defined increase in the maximum metabolic rate with incubation temperature (Vetter et al. 1983, Simcic et al. 2015, Sear et al. 2016). For example, in a similar study of steelhead, the maximum metabolic rate increased exponentially with incubation temperature as *R*_max_ = 17.3*e*^013*T*^ (Rombough 1986), which results in metabolic rates at MTDM of 72, 82 and 94 μgO_2_/h for 11, 12 and 13 °C respectively; a sufficient differential to demark a critical temperature at which the metabolic rate could exceed the oxygen supply for alevin. Thus, the steelhead study supports the possibility of temperature-dependent MTDM. However, laboratory studies with Sacramento winter-run Chinook salmon (USFWS 1999) do not support the hypothesis of alevin thermal mortality at *T*_*crit*_. Chinook eggs incubated as high as 13.3 °C experienced little mortality over development; increasing from 5% in egg cleavage, to 14% in embryos, and 17% in pre-emergent alevins. In contrast, at 15.5 °C pre-emergent alevin mortality increased from 17 to 78%, but remained unchanged in embryos. Thus, if the oxygen levels in redds were above the *P*_*c*_ for alevin it is likely they are largely protected from suffocation and mortality until temperatures approach 15 °C.

The temperature specific mortality rate *b*_*T*_ is closely linked with the duration of thermal susceptibility. Specifically, the ratio of *b*_*T*_ between models I and III reflects the ratio of the duration of thermal mortality susceptibility in the models. Because mortality, by definition, only occurs on a single day in models II and III, then the duration of susceptibility in model I is *b*_*T*_*III*__ / *b*_*T*_*I*__, which from Table 2 is between 80 and 100 d and therefore equivalent to the incubation duration. Thus, differences in *b*_*T*_ in the models are reconciled by the model assumptions on the duration of mortality. In short, *b*_*T*_ estimated in model I likely underestimates the effective field value and overestimated the duration of thermal susceptibility.

### Spatial dependence of carrying capacity model

Models I and II estimate carrying capacity directly as a free parameter. Thus, models I and II use the redd numbers over the entire habitat and calculate a single carrying capacity for the entire habitat. In contrast, model III uses the observed redd numbers for each reach and estimates the carrying capacity in terms of the number of redds per unit length of reach, i.e. *β*. Thus, the carrying capacity for each reach is calculated as *K*_*i*_ = *β*_*i*_*L*_*i*_ where *L*_*i*_ is the length or reach *i*. Models I and II each have 4 fitting parameters to fit and model III has 5 fitting parameters. However, in using reach-specific redd numbers model III contains more information than models I and II. Table 3 compares the carrying capacity estimates of the three models by converting redd counts to carcass counts using the factor 5.7 carcass detections per redd detection. Models I and III carrying capacity estimates were similar but model II carrying capacity was lower by a factor of 5 (Table 3). Importantly, the base survivals *S*_0_ were different, with model III survival 30% higher than in model I (Table 2).

**Table 3.** Model carrying capacities in carcasses for entire habitat *K* and for specific reaches *K*_*i*_ for model III using the formula *K*_*i*_ = 5.7*βL*_*i*_ where *L*_*i*_ is the reach length from Table 1. Few redds were ever observed in reach 5 so it is removed from the habitat calculation.

| Model | $K$ | $K_1$ | $K_2$ | $K_3$ | $K_4$ |
| --- | --- | --- | --- | --- | --- |
| I | 9107<sup>a</sup> | - | - | - | - |
| II | 1842 | - | - | - | - |
| III | 8561 | 1208 | 707 | 4024 | 2622 |
| III | 9160 | 1294 | 758 | 4298 | 2810 |

a. obtained from (Martin et al. 2017)

The partition of spawners by reach provides information on the spatial interactions of spawners with density and temperature. Between reaches 1 and 2 the fractions of the total redds exhibit an inverse relationship while the fractions in reaches 3 and 4 are uncorrelated (Fig. 3). The inverse correlation might involve contagion effects in which spawners were attracted to previously created redd sites (McNeil 1967, Essington et al. 1998). The lack of correlation between reaches 2 and 3 may involve the low fish densities in reach 3 compared to the densities in 1 and 2, i.e. the densities are too low to produce contagion behavior.

Finally, the partition of fish between reaches may also be influenced by the pre-spawning temperature (Strange 2010, Moore et al. 2012). Such effects have been observed elsewhere. For example, in Oregon, cold water patches are associated with increased abundance of salmonids (Ebersole et al. 2003) and studies indicate salmonids delay spawning at temperatures above 13 °C (Richter and Kolmes 2005). For the Sacramento River, the downstream increase in pre-spawning temperature (Fig. 2a) may limit the suitable spawning habitat. Figure 4, illustrating redd fractions in reaches against the pre-spawning temperature in reach 4, suggests a possible effect. The fraction of redds in reaches 1 and 2 are independent of reach 4 temperature (Fig. 4 a, b) but the fraction in reach 3 is below the across-year mean for reach 4 temperatures above 12 °C (Fig. 4c). This suggests that when downstream temperatures are warm, spawners bypass reach 3 and move into reaches 1 and 2 further upstream. Interestingly, the threshold for this hypothesized response corresponds with the critical temperature for the onset of egg thermal mortality, suggesting the prespawning response to temperature may have evolved to provide fitness to the offspring.

### Implications of the models to temperature management

The three models, using similar coefficients, reasonably fit the observed survival patterns of winter-run Chinook salmon between egg deposition and fry passage at RBDD. However, the implications of the models to temperature management are different and provide different answers to the question proposed by (Sabo et al. 2017) of “when is water most needed and when can it be spared”. Based on model I, temperature should be controlled between the beginning and end of the spawning season while based on model III the duration of control would extend between egg hatching dates for the first and last redds of the season. NMFS, developing a management plan based on model I, proposed controlling temperature between Keswick Dam and the confluence with Clear Creek (Fig. 1) from May 15 through October. Applying model III, and noting that hatching occurs about midway through the incubation, the same level of thermal protection might be obtained with temperature control between July and September; reducing the period of cold water releases from the reservoir from over five months to as short as three months.

Model III suggests the possibility of additional benefits in dryer years. Because the volume of reservoir cold water is more limited in dryer years, the compliance temperature might have to be raised or the duration of temperature control might have to be reduced. The strategy proposed by NMFS, based on model I, maintains the duration and raises the compliance temperature in dryer years. Under model III the duration of temperature control would be shorter allowing for a smaller increase in compliance temperature. In essence, model III scenarios might provide the same thermal protection as model I scenarios but use less water in normal years and better protection than model I in dry and critical dry years.

Model III also suggests additional management strategies that optimize the trade-off between thermal-and density-dependent mortalities. Firstly, as discussed above, controlling pre-spawning temperatures may reduce or expand the extent of the spawning distribution in advantageous ways. Under dry and critically dry conditions and low run sizes, the pre-spawning temperature might be elevated to encourage the adult population to spawn closer to Keswick Dam. This action might reduce thermal mortality without a significant increase in background mortality. Correspondingly, with an adequate cold water supply and above average run sizes, the pre-spawning temperatures might be lowered to expand the spawning distribution and thus reduce background mortality. By accounting for the effect of river temperature on age of hatching, an extension of model III could explore the possibility of accelerating the hatch time and thus reducing the period of thermal sensitivity of the population.

While model III uses the greatest amount of information on spawning distributions for estimating background effects, it only considers the mean spawning dates. Thus, it does not capture the range in spawning distributions over a season which can extend three months (B. Martin personal communication). However, restricting the analysis to the mean spawning date had practical benefits and was judged to be sufficient for two reasons. Firstly the temperature in most years was relatively constant over the spawning season so with a Gaussian spawning distribution the mean spawning date generally represents mean conditions experienced during incubation. The second reason the mean values were use was because the egg-to-fry survivals to which the model was fit are seasonal averages. Thus, it did not seem much would be gained by matching season-specific information on spawning with seasonally averaged survivals. Additional work is currently underway that addresses the seasonally specific spawning and survival information.

In summary, the models illustrated here highlight the importance of characterizing the biological and spatial/temporal details of fish early life growth and survival when developing temperature control programs for managed river systems. Characterizing and modeling biological responses for target species such as winter-run Chinook is essential for achieving efficient reservoir operations (Adams et al. 2017). In the larger context, the models illustrate the value of the emerging perspective of water management from simply meeting compliance goals to meeting and managing the biological needs of the species.

## Acknowledgement

This work was funded by the San Luis and Delta-Mendota Water Authority and Bureau of Reclamation contract R17AC00158. I wish to acknowledge S. Greene, C. Hanson and J. Israel for providing ideas and data during the development of the paper and R. Buchanan for assistance in developing the fitting algorithm.

---

**Figure S1.**
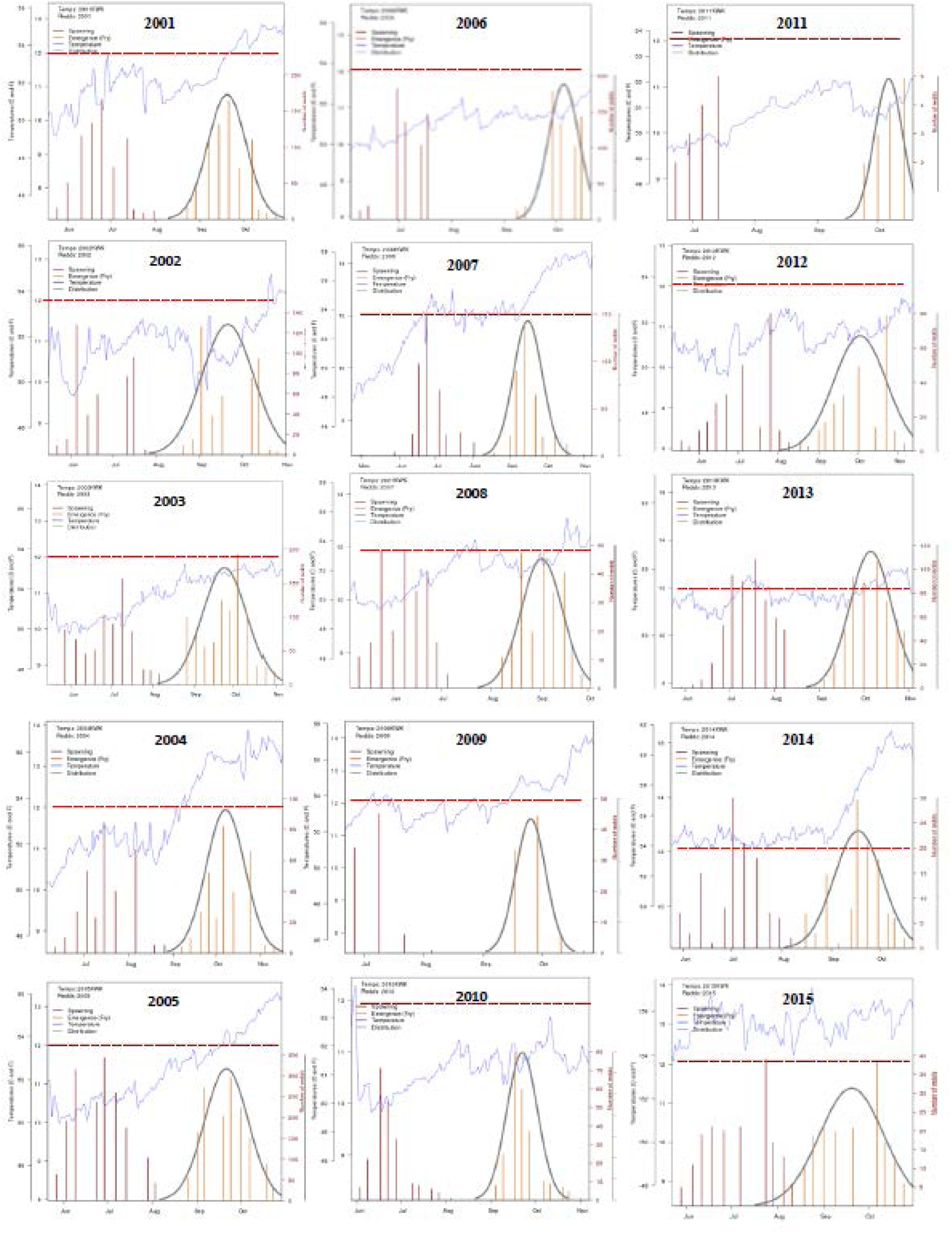
Temporal patterns of river temperature at Keswick Dam, redd distribution and predicted dates of fry emergence. Graphs were generated from the SacPAS model available at http://www.cbr.washington.edu/sacramento/migration/

